# Humans incorporate trial-to-trial working memory uncertainty into rewarded decisions

**DOI:** 10.1101/306225

**Authors:** Maija Honig, Wei Ji Ma, Daryl Fougnie

**Affiliations:** New York University Center for Neural Science & Department of Psychology; New York University Abu Dhabi Department of Psychology

## Abstract

In daily life, working memory plays an important role in action planning and decision-making. However, both the informational content of memory and how that information is used in decisions is poorly understood. To investigate this, we conducted a memory experiment where people not only reported an estimate of a remembered stimulus, but also made a rewarded decision designed to reflect memory uncertainty. Reported memory uncertainty is correlated with estimation error, showing people incorporate their trial-to-trial memory quality into rewarded decisions. Moreover, memory uncertainty can be combined with other sources of information; after we induced prior beliefs about stimuli probabilities, we found that estimates shifted towards more frequent colors, with the shift increasing with reported uncertainty. The data is best predicted by models where people incorporate their trial-to-trial memory uncertainty with potential rewards and prior beliefs, highlighting the importance of studying working memory as a process integrated with decision-making.

---

Working memory (WM), the storage and manipulation of information on a short timescale, is essential for many cognitive processes. For instance, individual differences in WM predict intelligence and academic success^1–3^. To understand WM, research has largely focused on examining the capacity and limitations of WM (reviewed in Luck and Vogel, 2013^4^, Ma et al., 2014^5^), epitomized by memory paradigms such as delayed estimation^6–8^, in which participants report a guess of a stimulus feature after a delay. In real life, however, WM information is not only used to make estimates of stimuli features, but to make decisions and take actions. For example, when deciding when to cross the street, a person must remember from a glance the position and velocity of cars. Since a mistake in this decision is costly, and memories are noisy, WM information ought to be combined with information about potential rewards (getting to your destination sooner) and costs (getting hit by a car) of the decision. In doing so, it is useful to know how reliable, or conversely how uncertain, one’s memory is. If someone is uncertain about the speed of an oncoming car, they might play it safe and wait a little longer in order to avoid a high-cost collision.

Yet uncertainty in working memory is rarely studied. There is evidence that people know something about the quality of their WM representations^9–12^. For example, people can report which items from a set of stimuli they remember better^9^. Beyond this, people may have knowledge of their uncertainty on a trial-to-trial level: when people make explicit reports of confidence in memory decisions, the amount of response error correlates with the reported confidence on each trial^10;12^. This could be explained by memory confidence being a function of internal fluctuations in underlying memory quality^13^. However, confidence ratings are not necessarily a reflection of memory uncertainty^14;15^ and may be produced through a different mechanism from those used to make decisions under uncertainty^16;17^.Thus, to observe memory uncertainty, experimenters may not want to rely on *explicit* reports of uncertainty alone, but also use paradigms in which people are incentivized to make decisions which *implicitly* incorporate memory uncertainty by combining it with other information.

To benefit from storing trial-to-trial information about memory quality, an observer must not only know their memory quality, but be able to combine it with other information to make a decision. For example, when crossing the street, if it is rush hour when cars are likely to appear quickly, an observer ought to have a higher standard of certainty that a car is far away before crossing. This is the case in perception, where people are able to improve their decisions by incorporating their uncertainty with other sensory information^18;19^, rewards^20–23^, or prior beliefs^24;25^. However, though there may be a close connection between perceptual and working memory representations^26–28^, much less is known about the role of WM uncertainty in memory-based decisions. People combine WM reliability and peripheral sensory information to plan reaching movements^29^ and humans and monkeys perform near-optimally in change detection tasks when items vary in reliability^30;31^. However, both in perception and memory, these studies typically manipulate uncertainty through varying the encoding precision of stimuli (and thus the resulting memory uncertainty). This is done by manipulating the reliability of stimuli, by varying properties such as contrast, density, or visual eccentricity, creating the possibility that participants are using the visible reliability of stimuli as a proxy for memory uncertainty^32^. In this way, participants could use cues about such attributes as a stand in for memory uncertainty, without implicitly representing uncertainty about their memories. We aim to show that even in the absence of explicit uncertainty manipulations, WM represents a trial-to-trial measure of uncertainty which is taken into account in decisions.

We use a novel color working memory task to obtain on each trial not only a memory report, but also a continuous measure of trial-to-trial memory uncertainty through a rewarded decision. Furthermore, we manipulate the stimulus distribution to create expectations about likely stimulus values. In order to obtain rewards, participants must combine their memory uncertainty with information about the reward rule and the stimulus distribution. Our results show that WM represents uncertainty information, which people can combine with prior information and incorporate into rewarded decisions. This highlights the potential complexity of WM representations, and shows that rewarded decisions can be used as a powerful tool to examine WM and inform and constrain theoretical, computational, and neurobiological models of memory.

## Results

To explore how WM information is taken into account in decisions, we modify a delayed estimation task^6–8^ to include a rewarded decision, designed to reflect memory uncertainty. During each trial, participants viewed four colored circles and after a delay reported an estimate of the color of a probed circle. After responding, they were prompted to draw a symmetric “arc” around their estimate (Figure 1a, similar to^20^). We refer to this reported arc as the “arc size”, measured as half of the symmetric reported arc. If the true stimulus color was within the bounds of the arc (“hit”), participants received points which linearly decreased (100-0) as a function of arc size (Figure 1a); otherwise they received zero points (“miss”). This rule creates a trade-off between the probability of getting reward (as the arc gets bigger the stimulus is more likely to be inside it), and the magnitude of reward (as the arc gets bigger, points from a hit decrease). Participants were given monetary and time rewards based on points obtained, and were thus incentivized to use their memory uncertainty when reporting an arc (see Methods).

**Figure 1:**
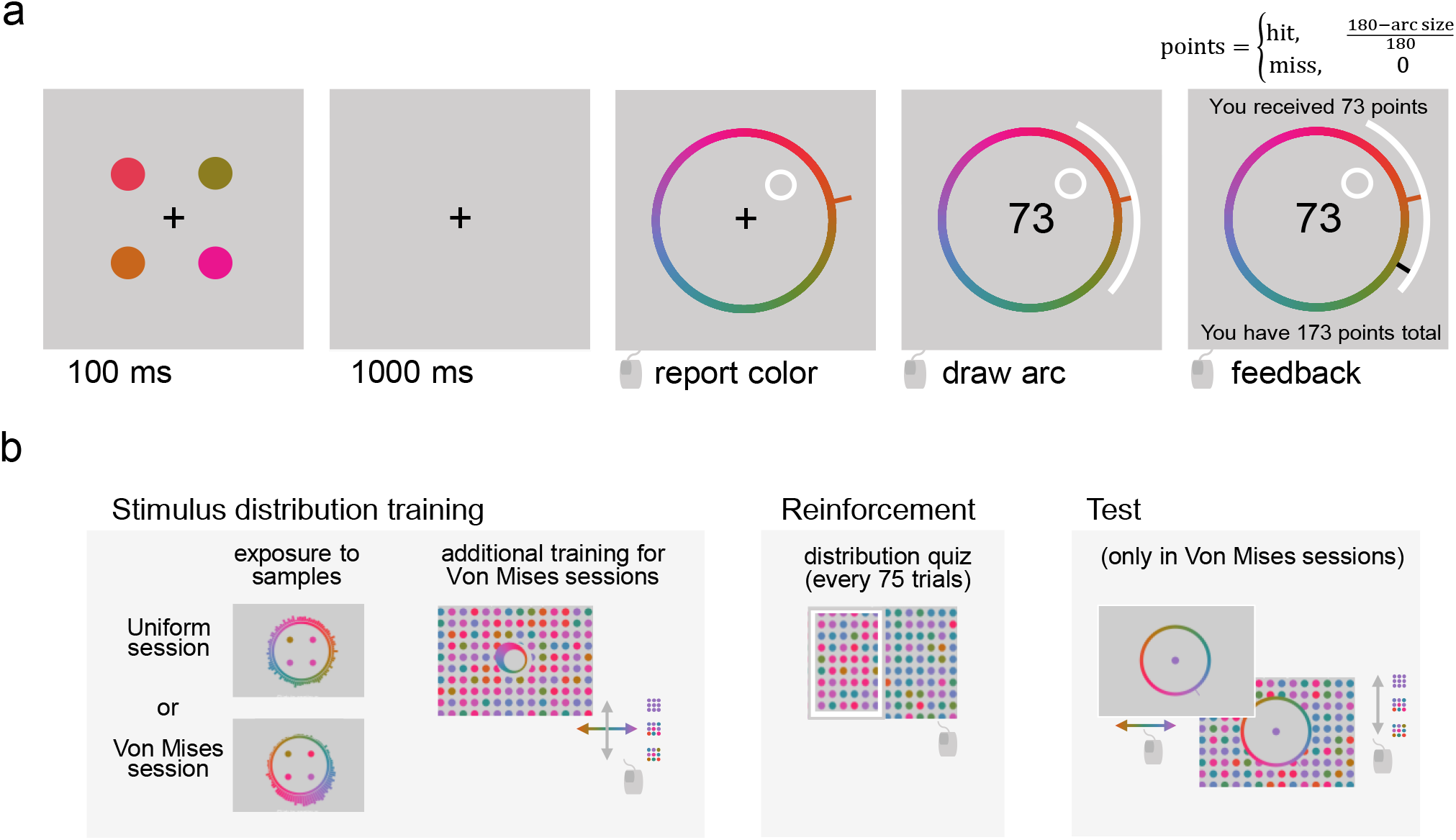
Experimental Design. (**a**) Trial procedure. Participants viewed four colored stimuli. After a delay, they estimated the color of a probed stimulus on a color wheel and then drew a symmetric arc around their estimate. If the stimulus value fell outside of the arc, the participant received 0 points (“miss”). If it fell within (“hit”), the number of points received was a decreasing linear function of arc size. (**b**) Stimulus distribution training, reinforcement, and testing. In a given session, the color distribution was either Uniform or von Mises. Each session started with exposure to 1000 *training* samples from the distribution. These samples were used to build a histogram representing the session’s stimulus distribution. In von Mises sessions only, the participant performed additional training by generating sets of dots matching the mean and width of the distribution (with feedback). After every 75 trials, the participant’s knowledge of the stimulus distribution was *reinforced* with a quiz. Participants had to identify the correct distribution from samples in a set of 2AFC questions (with feedback). At the end of each von Mises session, participant’s knowledge of the stimulus distribution was tested by having them report estimates of the mean and width of the distribution three times (by generating an example distribution of stimulion the screen).

Furthermore, to test whether working memory uncertainty could be integrated with other knowledge, we introduced expectations about stimuli colors (Figure 1b). In two of the four experimental sessions, stimuli colors were drawn from a uniform stimulus distribution, while in the other two, colors were drawn from a von Mises (circular normal) distribution. Participants were taught this distribution through extensive training (Figure 1b, see Methods). If prior and memory information are combined, we expect reponses would shift towards expected colors, particularly when participants were more uncertain.

To build and validate a model of WM uncertainty without having to account for prior beliefs, we first examined only the sessions where the stimulus distribution was uniform. Participants produced errors (defined as the circular distance between stimulus and estimate, in degrees) and arc sizes which follow smooth continuous distributions (Figure 2a-b). Across participants there were large variations in the distributions of arc sizes, possibly due to differences in risk attitude (see Supplementary material). Arc size is positively correlated with absolute estimation error on a single-trial level (“error-arc size correlation”, mean and s.e.m. across participants *r*_s_ = 0.31 ± 0.029, *t*(11) = 10.2, *p* = 6.1 · 10^−7^, Figure 2c), such that riskier decisions (smaller arcs) were associated with higher errors and vice versa. Thus, people have knowledge of their own memory quality, consistent with the idea that WM stores information about memory uncertainty. Furthermore, across participants the ability to report arcs that optimize awarded points (see Supplementary material), was inversely correlated with absolute estimation error across participants (*r_s_* = 0.80, *p* = 0.0032), suggesting that memory quality and the ability to know one’s memory quality are related.

**Figure 2:**
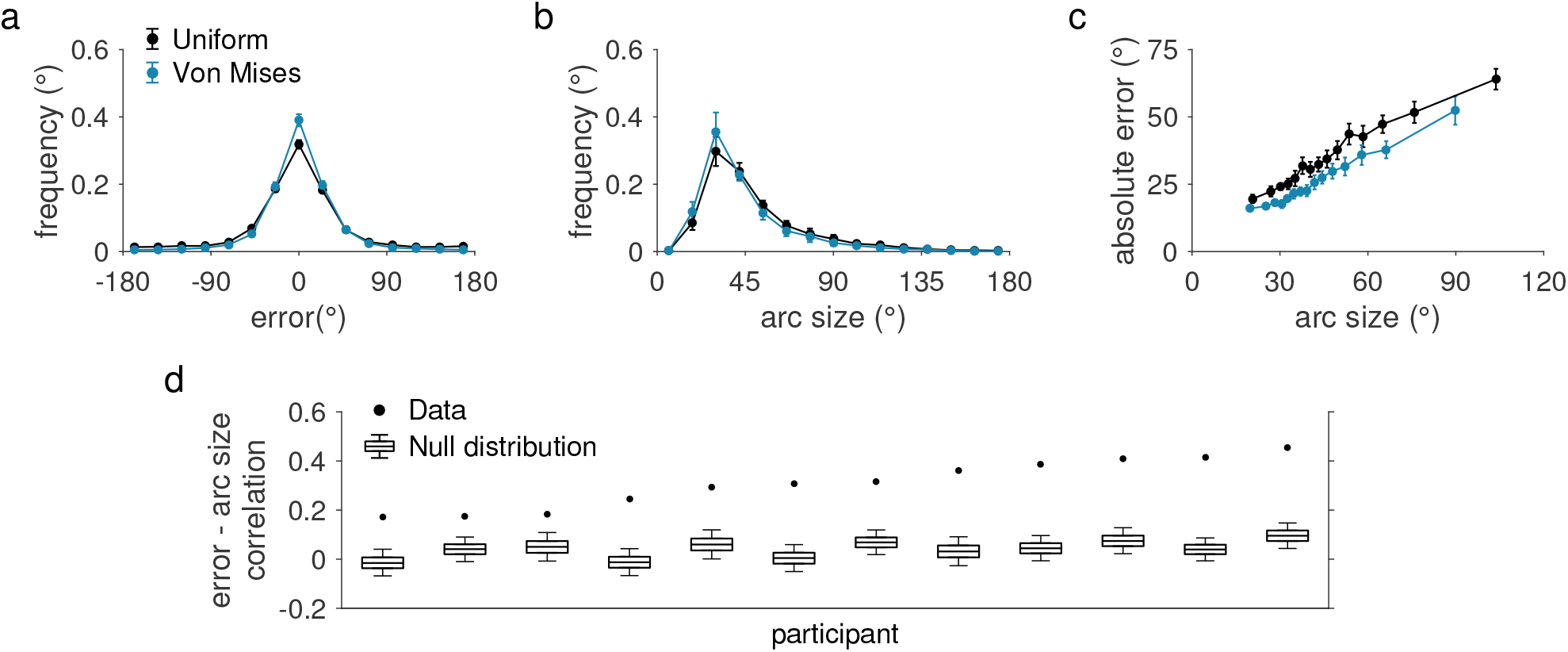
Arc size reports correlate with error and reflect trial-to-trial knowledge of memory quality. Behavioral data from uniform stimulus distribution trials (black) and von Mises stimulus distribution trials (blue). Here and elsewhere, error bars represent mean and standard error of the mean across participants. (**a**) Histograms of circular estimation error in 24° bins. (**b**) Histograms of arc size in 12° bins. “Arc size” refers to half of the size of the reported arc. (**c**) Error and arc size, plotted in 15 quantile bins. For each participant we separate the data into 15 quantiles of arc size and calculated the mean arc size and absolute estimation error per quantile. For each quantile we then calculated the mean and s.e.m. of these averages across participants and plotted the point at the horizontal coordinate equal to the mean of the quantile centers. Error is correlated with arc size, suggesting trial-to-trial knowledge of memory quality. (**d**) Comparison of the observed error-arc size correlation coefficient to the null distribution expected if the correlation were solely driven by stimulus color (see Methods). Box plot: 5th, 25th, 50th, 75th, and 95th percentiles. The observed correlation values fall far outside of the null distribution, indicating that they cannot be solely driven by stimulus color.

However, being able to report about memory quality does not necessarily imply that memory stores a representation of memory uncertainty. Observers can have indirect knowledge of memory quality if they rely on factors that correlate with their internal noise, such as knowing which stimuli are most difficult to encode, or knowing their own attentional state. To show that the observed relationship between error and arc size reflects knowledge of memory uncertainty, we rule out several other interpretations. One possibility is that people use information about stimulus features as a proxy for uncertainty. For instance, if an observer knows they encode the color pink poorly, they may use this knowledge to set a larger arc. To test this, we check if the observed error-arc size correlation can be explained by both variables being correlated with the stimulus color. We simulate error-arc size correlations from the null hypothesis (that correlation is caused by stimulus color, see Methods). If arc size solely reflects the influence of stimulus color, and not internal memory uncertainty, then we would expect equally strong correlations between error and arc size in the simulated data. By contrast, we find that the observed correlations in the data (*r*_s_ = 0.31 ± 0.029) are far larger than the mean simulated correlations (Figure 2d, mean and s.e.m. *r*_null_ = 0.010 ± 0.040, *p* < 10^−9^ for all participants). We repeated this analysis for two other potential confounding factors: the dispersion of the colors within a display (mean and s.e.m. *r*_null_ = 0.016 ± 0.0068, *p* < 10^−9^ for all participants)^33^ and the distance in color space from a probed color to the nearest non-probed item (mean and s.e.m. *r*_null_ = 0.0063 ± 0.0082, *p* < 10^−9^ for all participants)^34^, both of which predict mean correlations much lower than the observed correlations. Furthermore, our data shows little evidence for color biases^35;36^ or dependencies between trials^37;38^ which could also contribute to knowledge of encoding precision (see Supplementary Material). These results suggest that the correlation between error and arc size does not arise solely from participants representing stimulus-related qualities as a proxy for memory uncertainty.

To further confirm that arc sizes are a reflection of memory uncertainty, we build a computational model that jointly predicts errors and arc sizes. We assume that stimuli are encoded as noisy memories (represented by a von Mises distribution), with an encoding precision that varies per trial (variable-precision/VP model) according to a gamma distribution (2 free parameters)^9;39;40^. We make the additional assumption that the observer stores a representation of their encoding precision as memory uncertainty, the width of the likelihood, which is combined with prior beliefs to generate a posterior distribution over stimuli colors. To obtain an estimate, we assume that observers sample from a noisy representation of the posterior (free parameter)^41^. To obtain an arc size, we assume the observer first calculates the expected utility of each possible arc response by combining reward and memory uncertainty information, and then samples from a noisy representation of expected utility (softmax noise, free parameter) (Figure 3). To account for different risk attitudes across individuals, we allow for a power law nonlinearity (free parameter) between reward and utility^42^. Finally, we assume that on some proportion of trials (free parameter) the observer fails to encode the stimuli, and responds as if they had no information (lapse trial). We fit this model and compare it to several alternative models using the summed 10-fold cross-validated log-likelihood (LLcv) to account for differences in model complexity (see Methods). Tables of fitted model parameters and model predictions are available in the Supplementary material. The resulting model (**VP-known**, 6 parameters) describes human error-arc size distributions qualitatively well (Figure 4a-c). Removing the utility non-linearity (*α*), the decision noise (*ν*), or lapse rate (*λ*) significantly worsens the fit (see Supplementary material). Parameters and model comparison results are consistent across stimulus distributions, and consistent when fit only on the stimulus estimate (analogous to delayed estimation, see Supplementary material).

**Figure 3:**
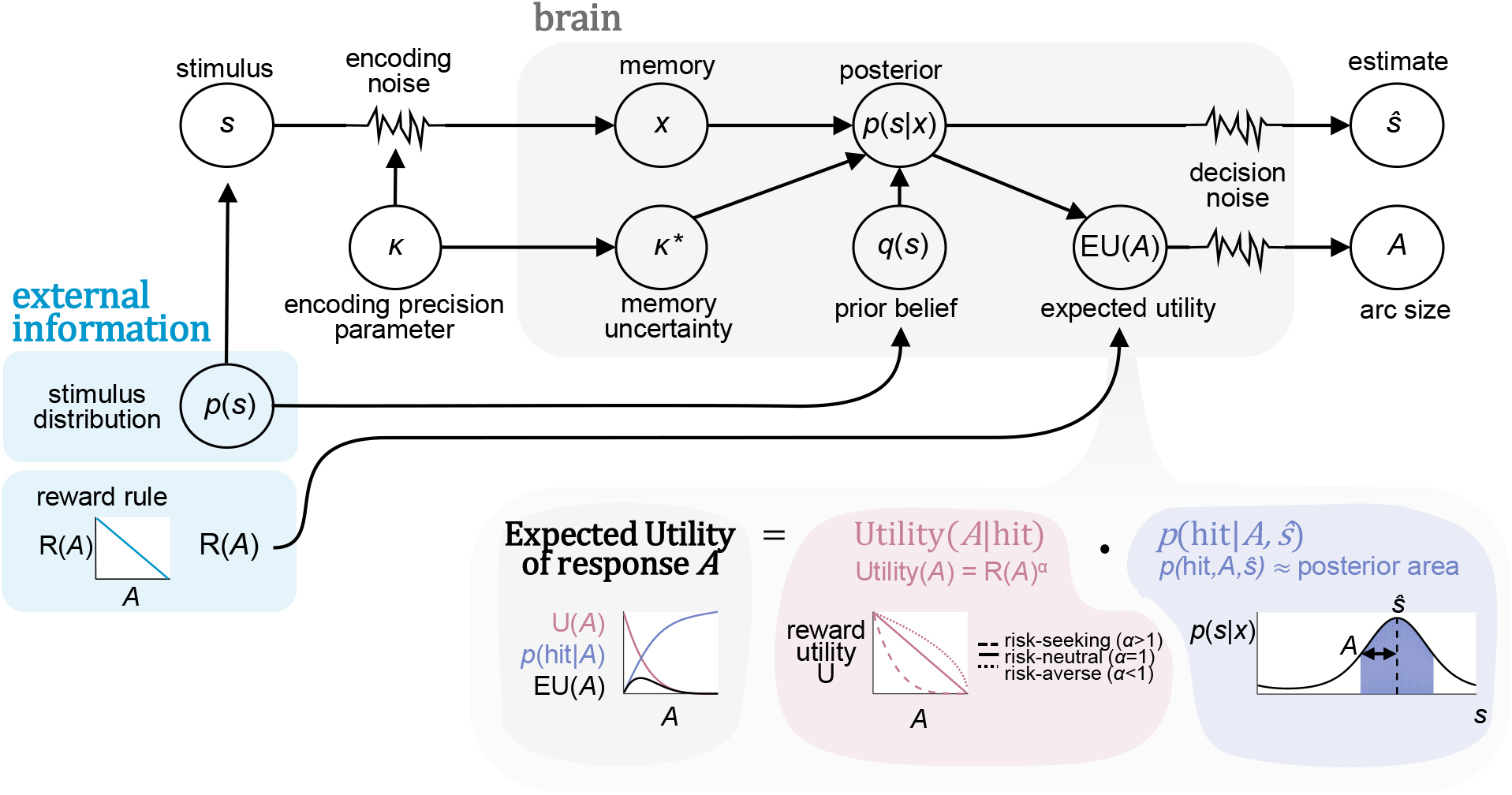
Schematic of the computational model. The observer’s memory *x*, of a color *s*, is noisy, parameterized by a von Mises distribution with encoding precision parameter *κ* (the concentration). Memory uncertainty is the width of the likelihood function (reflecting observer’s beliefs) over the remembered color, represented by *κ*\*, which is a function of *κ*. An ideal observer would combine their likelihood *p*(*x*|*s*) with prior beliefs *q*(*s*) to obtain a posterior *p*(*s*|*x*). The stimulus estimate 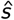 is sampled from the posterior distribution raised to a power (representing decision noise). The observer computes the expected utility of potential arc sizes *A* by multiplying their utility *U* given a hit with the probability of a hit under the posterior distribution. To obtain an arc size the observer samples from a noisy version of the expected utility (softmax). The utility of *A* is the number of points earned raised to a power α to capture risk attitude.

**Figure 4:**
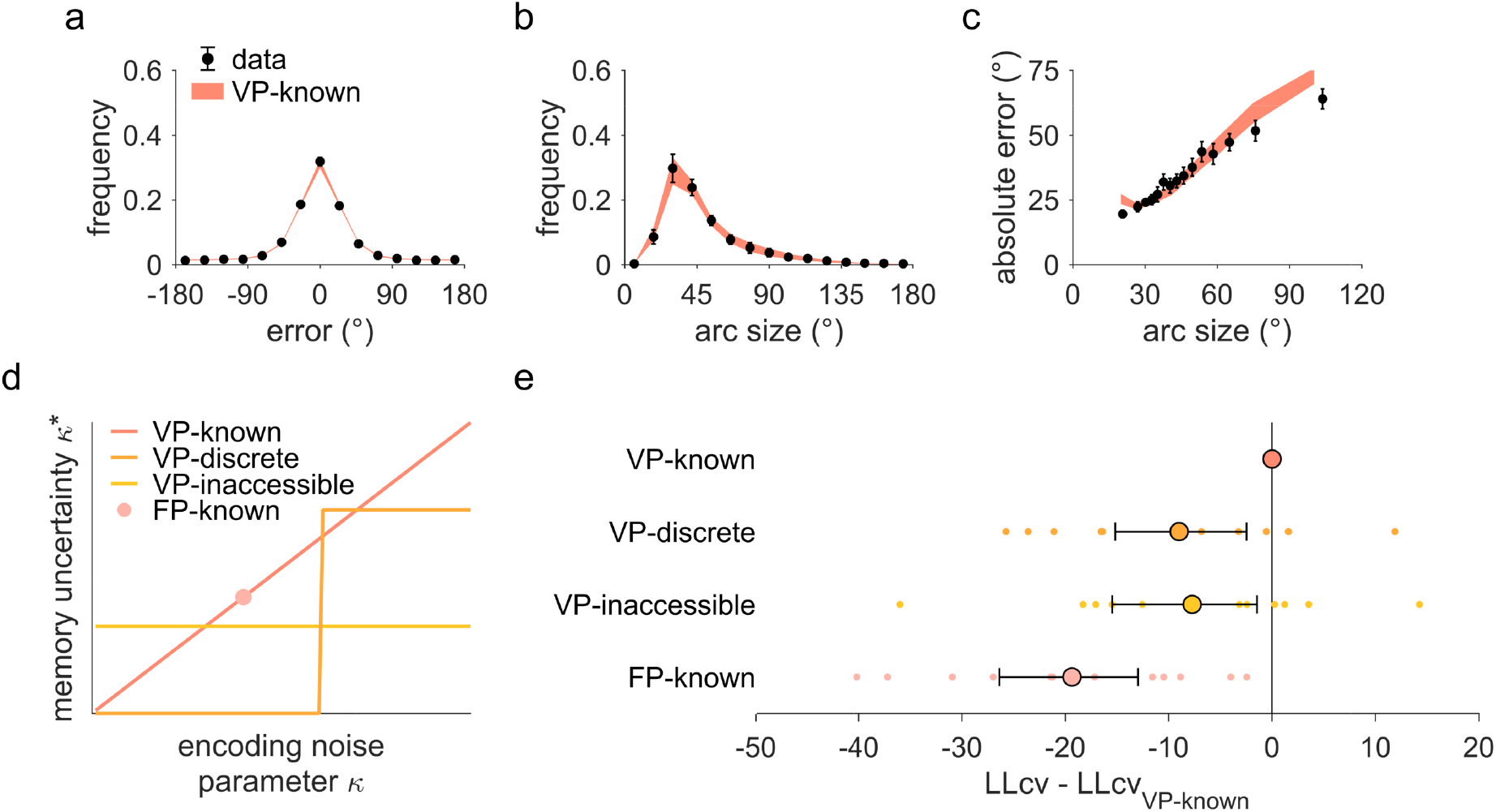
Model comparison based on sessions with a uniform stimulus distribution. (**a-c**) VP-known model predictions, plotted with the same conventions as Figure 2. In this model, observers encode stimuli with variable precision and have full knowledge of their encoding precision *κ* as memory uncertainty *κ*\*, (*κ*\* = *κ*). (**d**) Schematic representation of the relationship between *κ*\* and *κ* in the models tested. VP-known: observers know their uncertainty perfectly. FP-known: observers encode stimuli with a single fixed precision level, which they know and use to report an arc. VP-inaccessible: variable-precision encoding, but observers have no access to *κ*, and instead assume a fixed memory uncertainty *κ*_assumed_ (free parameter). VP-discrete: variable precision encoding, but observers have limited knowledge of *κ* and represent it as either 0 (when below a threshold *κ*_threshold_) or as a fixed value *κ*_assumed_ (both free parameters). (**e**) Model comparison using 10-fold cross-validated log-likelihood (LLcv). Dots: individual-subject differences between each model and the VP-known model. Circles and error bars: mean and 95% bootstrapped confidence interval. Negative numbers represent a worse fit than the VP-known model. The VP-known model fits the best, results are consistent when using AIC or BIC (see Supplementary material).

To examine what elements of this model are necessary, we modify it in several ways. First, we test a model in which encoding precision does not vary across trials, and is always fixed (**FP-known**, 5 parameters). Although encoding precision is fixed, an error-arc size correlation is sill predicted due to trials being a mixture of fixed-precision trials (small errors and arcs) and zero-precision lapse trials (large errors and arcs). Model comparison results here and following are reported as the mean 10-fold cross validated log-likelihood difference from the best fitting model across participants (Δ_LLcv_), followed by a bootstrapped 95% bootstrapped confidence interval in brackets (see Methods). Critically, the VP-known model performs better than the FP-known model (Δ_LLcv_ = −19.2, [−26.3 – 12.9], Figure 4b). This suggests that memory quality is variable across trials, and that the error-arc size correlation is not entirely explained by an observer whose arc sizes are a function of their attentional state (lapse trials).

Even if encoding precision varies across trials, it is possible that observers do not have access to information about memory uncertainty when setting an arc. We test two models in which the participant has incorrect beliefs about their uncertainty. For instance, although stimuli are encoded with variable precision, an observer might not take into account their memory uncertainty, but instead always assume memory uncertainty is a fixed value (free parameter) when setting an arc (**VP-inaccessible**, 7 parameters). In this case, the observer still has some knowledge of memory quality: they are aware of lapse trials (in which they failed to encode the stimulus), on which they have higher error and report a larger arc size. Another possible scenario is that observers have only coarse, discrete knowledge of memory uncertainty (**VP-discrete**, 8 parameters). In this case, the observer sets an arc size with either a precision of zero or a fixed precision (free parameter) according to whether their encoding noise was above or below a certain threshold (free parameter) (Figure 4d). Although all models are capable of predicting a correlation between error and arc size, the VP-known model outperforms the VP-inaccessible (Δ_LLcv_ = −7.8, [−15.1 – 1.6]) and VP-discrete models (Δ_LLcv_ = −9.1, [−15.7 – 2.7], Figure 4c). This suggests that participants incorporate their uncertainty into arc size reports and suggests that WM uncertainty is a causal factor of both estimation errors and arc responses.

To test whether people could combine their working memory uncertainty with prior information, we manipulated the stimulus distribution: in half the sessions stimuli were generated from a von Mises (circular normal) distribution, with a fixed width and a different mean for each participant. We trained participants to learn this distribution, inducing prior beliefs about stimuli (see Methods). Incorporating these prior beliefs with memory uncertainty would confer a performance advantage (see Supplementary material). For instance, if the color remembered is both highly uncertain and improbable given the believed stimulus distribution, a more accurate estimate can be made by shifting the response towards more frequent colors.

On von Mises stimulus distribution trials, absolute estimation error and arc size are significantly correlated (mean and s.e.m. *r*_s_ = 0.30 ± 0.029, *t*(11) = 9.8, *p* = 8.9 · 10^−7^, Figure 2c), consistent with previous findings that people have knowledge of their memory quality^9;10;12^. Additionally, in von Mises stimulus distribution trials there is less error compared to uniform stimulus distribution trials (mean error_Unif._ = 36° ± 1.9°, mean error_VM_ = 27° ± 1.8°, Wilcoxon signed-rank test *p* = 4.8 · 10^−4^) and lower reported arc sizes (mean arc_Unif._ = 49° ± 4.1°, mean arc_VM_ = 44° ± 3.9°, Wilcoxon signed-rank test *p* = 0.0015), consistent with the idea that participants use prior beliefs to lower their uncertainty and make more accurate estimates (Figure 2a-b). If people incorporated prior information into their stimulus estimates, their responses should be attracted towards the most frequent color. Validating this, responses on the left (‒) of the most frequent color (0) have rightwards error (+) and vice-versa (Figure 5a). We quantify this by multiplying the circular error by the sign of the stimulus on that trial to obtain the directional shift towards (+) or away from (‒) the prior. The shift towards the prior is positive, showing that stimulus estimates are attracted to the prior mean (mean shift = 3.1° ± 1.38, Wilcoxon signed-rank test *p* = 0.034). Importantly, the amount of shift towards the prior mean increases with increasing reported arc size, suggesting that people incorporate prior and memory information in proportion with their memory uncertainty (mean *r*_s_ = 0.080 ± 0.025, *t*(11) = 3.3, *p* = 0.0077, Figure 5b).

**Figure 5:**
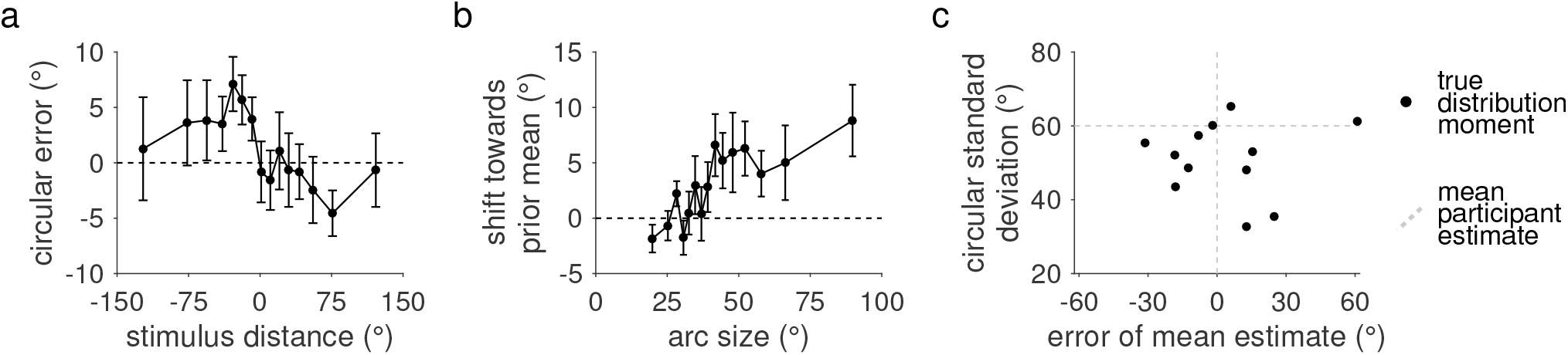
Prior information is learned and incorporated into decisions. Behavioral results from sessions with a von Mises stimulus distribution. Plots are binned in 15 quantiles of either stimulus distance from the prior or arc size as in Figure 2c. (**a**) Trial-averaged circular estimation error against stimulus distance from prior mean. For stimuli counterclockwise (‒) of the prior mean, errors were in the clockwise direction (+) and vice versa, showing that responses were shifted towards the prior mean (0). (**b**) We define “shift towards the prior mean” as the trial-averaged error either towards (+) or away from (‒) the prior mean. As the reported arc size increases, responses shift more towards the prior mean. This suggests that memory uncertainty modulates the influence of prior information in the decision. (**c**) Scatterplot of the mean estimates of the moments of the stimulus distribution, explicitly reported six times at the end of the experiment (Figure 1b). Plotted is the mean distribution mean estimate, and the mean circular standard deviation estimate across the six guesses. Responses indicate that some participants were fairly accurate at reporting the prior mean while others were very inaccurate (mean error > 30 degrees on a 360 degree color wheel). Participants also chronically underestimated the width of the prior distribution.

This suggests that when combining memory and prior information, people are able to weigh their memory information in proportion to their memory uncertainty. To understand how people do this, we take the model that best explains the uniform stimulus distribution data (VP-known), and add prior information to it, jointly fitting the model on uniform and von Mises stimulus distribution data. However, introducing a prior to the VP-known model introduces two issues. First, how does the observer combine prior and memory uncertainty? Second, what are an observer’s prior beliefs about the stimulus, and have they learned the stimulus distribution that was trained?

An optimal observer would incorporate prior information into their decision. Specifically, they would use Bayes’ rule to combine the likelihood (and associated memory uncertainty) and the prior beliefs to obtain a posterior. However, observers may just ignore prior information. Furthermore, although participants were trained until they learned the stimulus distribution (see Methods), they may not have learned the stimulus distribution correctly and may have incorrect internal beliefs about the prior.

To test whether participants learned the stimulus distribution, we assessed their knowledge of the distribution at the end of each von Mises session by having them report three estimates of the distribution moments (mean and circular standard deviation, for a total of six guesses across the two sessions, see Methods). Overall, participants were fairly accurate in estimating the mean (mean and s.e.m. absolute error= 18° ± 4.5) of the stimulus distribution, but underestimated the circular standard deviation (mean and s.e.m.=‒10° ± 2.5, Wilcoxon signed-rank test *p* = 0.015). Furthermore two participants performed particularly poorly (mean error > 30 degrees), suggesting that not all participants learned the stimulus distribution. To account for these deviations, our models allow for the possibility of observers incorrectly learning the width of the prior distribution. We we let the prior concentration parameter be a free parameter, *κ_w_*, which is fitted to the individual participant behavior (“Yes prior model”, **YY**). To validate this choice, we also test a special (nested) case of the YY model, in which the observer uses the true prior width (“True prior model”, **TT**, *κ_w_*=1.422). The TT model fits worse than its parent model YY (Δ_LLcv_ = ‒25.3, [‒31.9 – 16.4]), suggesting that participants may have incorrectly learned or represented prior information.

We compare the YY model to a model in which observers do not use prior information in any way (“No prior model”, **NN**). The YY model outperforms the NN model (Δ_LLcv_ = ‒15.1, [‒21.7 – 10.0], Figure 6b), suggesting that participant responses incorporate prior information.

**Figure 6:**
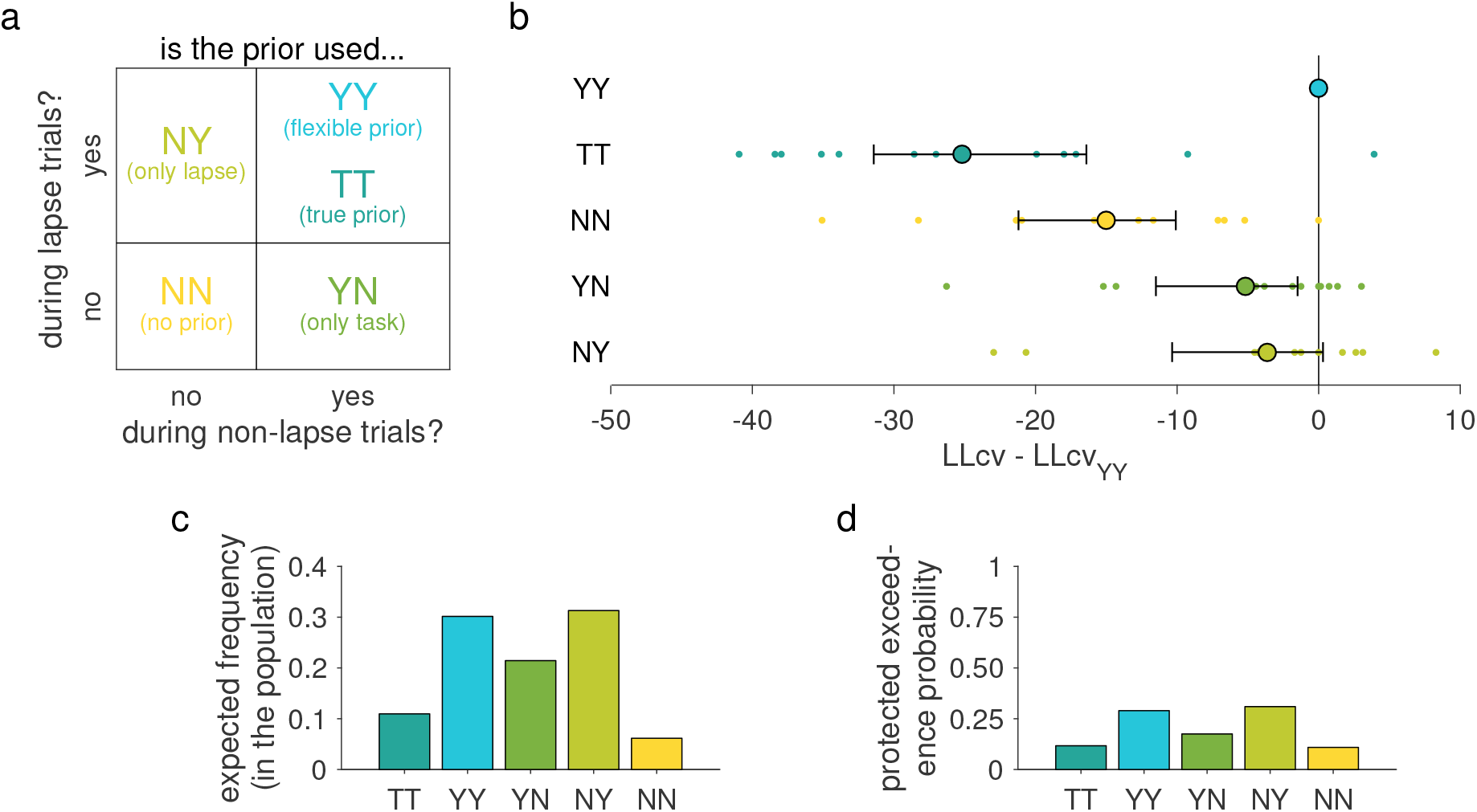
Comparison of prior use models. Comparison of five models that differ in the way by which prior information is used. (**a**) We define each prior-use strategy by whether (“Y”) or not (“N”) observers use the prior during non-lapse (first Y or N) and lapse trials (second Y or N). A special case of the YY strategy is one in which the observers use the true stimulus distribution as their prior (TT). (**b**) Model comparison using summed 10-fold cross-validated log-likelihood (LLcv). Dots: individual-participant differences between each model and the YY model. Circles and error bars: mean and 95 % confidence interval. Negative numbers represent a worse fit than the YY model. Models which ignore the prior (NN) or use the true prior width (TT) perform poorly. The YY model describes the data best overall, but does not describe each participant best. Results are consistent when using AIC or BIC (see Supplementary material). (**c**) Expected frequencies of the models in the population, obtained using Bayesian Model Selection. Overall, most participants are expected to use the prior in some way (YY,YN,NY,TT), but are split between participants who combine the prior with memory uncertainty (YY,YN,TT) and those who do not (NY). This suggests that the way that individuals use prior information in this task varies. (**d**) Protected exceedance probabilities of model obtained from Bayesian Model Selection. This is the probability that a given model is the most likely model in the population, which is low even for the most likely model (NY).

The YY model also captures the qualitative trends of the data (see Supplemental material), including the correlation between response shift towards the prior and arc size. However, this correlation may not result from the combination of prior information and memory uncertainty, but from an observer who uses prior information only when highly uncertain (i.e. a lapse trial). This would generate a mixture of high uncertainty high shift trials and low-uncertainty zero shift trials. To test this, we test models in which participants exhibit different prior-use behavior in lapse trials compared to non-lapse trials. On a lapse trial, where the observer has no information about the stimulus, a prior-using observer would respond at the mean of the prior (since the likelihood distribution is flat), while a non prior-using observer would respond with a random (uniform) guess when lapsing. We denote models with two letters, the first indicates whether prior information is used (“Y”=yes, “N”=no) during non-lapse trials, while the second indicates whether prior information is used while lapsing (Figure 6a). In this framework the YY and NN models are the same as above (respectively using, and not using prior information). However, we introduce a new model, in which the observer only uses prior information when lapsing (**NY**). This would generate a mixture of non-lapse trials with no prior shift, and lapse trials with a large amount of prior shift, creating a correlation between prior shift and arc size. For completeness, we also test the opposite model, where participants use prior information during non-lapse trials, but respond from a flat likelihood distribution when lapsing (**YN**).

We equipped the two new models (YN, NY) with a free parameter representing the concentration parameter of the prior (as in the YY model), and fit them jointly on both uniform and von Mises stimulus distribution trials. While the YY model still has the most predictive power, we find that both the NY (Δ_LLcv_ = ‒3.4, [‒10.2 0.6]) and YN (Δ_LLcv_ = ‒5.5, [‒12.4–1.6]) models fit the data almost as well (Figure 6b). However, despite small mean differences in model evidence, there is high variance in model evidence differences across participants. Even the model with the most predictive power (YY) does not describe many participants best, suggesting that people could have different strategies of prior use, better explained by different models.

To examine whether different people could have different strategies of using prior information, we assume that our participants do not all use the same model, and examine an alternative, hierarchical, model of how participants could be distributed using Bayesian Model Selection (BMS)^43^. This is a hierarchical model which treats models as random variables (“random effects”) and estimates the parameters of a Dirichlet distribution describing the probability of each model given the log-evidences (in our case the summed 10-fold cross validated log-likelihood) for each model and participant. By applying this hierarchical model we can obtain those Dirichlet parameters and thus the probability of each model occurring in our participant population (Figure 6c), as well as the protected exceedance probability, the probability that a given model is the most likely in the population above and beyond chance^44^. Overall most (62 %) of participants are estimated to come from models in which they use the prior with memory information (YY: 0.29, YN: 0.22, TT: 0.11) while the other participants either do not use the prior (NN: 0.06) or use the prior without memory uncertainty (NY: 0.32). The protected exceedance probability is low even for the maximally represented model NY (0.31, Figure 6d, see Supplementary materials), suggesting there is no dominating model among our participants.

This suggests that people can incorporate prior information and memory uncertainty in simple decisions, though not all people may do so. To confirm that participants use different strategies, we compare the probability of participants all being from the same model (“fixed effects hypothesis”) to the probability of participants being from different models (“random effects hypothesis”, see Methods). The random effects hypothesis is far more probable than fixed effects (random/fixed odds-ratio = 5.74 · 10^4^), suggesting that it is more likely that strategies of prior use vary across individuals, then that they use any single strategy captured by our models. Regardless of the underlying algorithm, we suggest that individuals can combine prior information with memory uncertainty. Further work will be needed to understand how prior information and memory uncertainty are combined.

## Discussion

Working memory is crucial to many aspects of human cognition and behavior, yet we do not fully understand what information is stored in WM or how it is incorporated into decisions. We examined this using a visual working memory task where people reported memory uncertainty through an incentivized rewarded-decision. Our measure of memory uncertainty, arc size, is highly correlated with performance, demonstrating that people know something about the quality of their own memories. This is consistent with previous work showing that people can report about their memory quality^9–11^. We further show that people’s knowledge of memory quality cannot be explained by knowledge of memory quality stemming from either stimulus display factors or lapse trials, suggesting that WM represents memory uncertainty. Our modeling suggests that spontaneous fluctuations in this memory uncertainty are the common cause of both estimation errors and arc sizes in this task.

Furthermore, memory uncertainty can be incorporated with prior beliefs about the environment. We manipulated people’s expectations about stimuli colors by changing the stimulus distribution so that some colors appeared more often than others. When participants believed that the color distribution was nonuniform, reports were attracted towards the most frequent color. Importantly, we found that the shift towards the prior increased with increasing memory uncertainty, consistent with the idea that people incorporate their memory information into decisions in proportion to their memory uncertainty^29;30;45^. However there may be individual differences in the way people combine their memory uncertainty and prior information. The data is best described by a hierarchical model in which different individuals use different strategies: some people use the prior with memory uncertainty in a Bayesian fashion, while others completely ignore the prior or use a different strategy. Future work will be necessary to understand the potential strategies and computations people use to combine prior and memory information; however, we have provided evidence that people are at least capable of such combination.

To draw this conclusion, we rely partially on models which assume that memory and prior information are combined with Bayes’ rule (YY, TT, YN, NY). However it is possible that people may have some other algorithm of combining information such as a heuristic strategy or weighted combination of memory and prior information. For instance, observers could combine prior information with a function other than Bayes’ rule. They could also could use a strategy where they switch between using the likelihood and the prior depending on their memory uncertainty (i.e. Laquitaine and Gardner^46^, this is similar to the NY model, which only uses the prior during lapse trials). While we do not examine other ways in which prior and memory information are combined, we hope that Bayesian combination with a flexible prior width provides a flexible framework where it is clear that individuals can combine prior information with memory information.

While we present evidence that WM contains a representation of memory uncertainty, it is an open question how this is implemented in the brain. In our models we assume memories are probabilistic representations similar to von Mises distributions, yet there are many ways probabilistic information could be represented in WM. The implementation most similar to our model is probabilistic population coding, where full continuous probability distributions over feature space could be represented in a population of neurons^47–49^. Though there is some evidence for this^50–52^, it is possible that a more limited form of probabilistic information is stored, such as a several samples of a stimulus, from which a memory uncertainty can be inferred^53–55^ or a scalar estimate of memory uncertainty^56^. We show that some kind of WM uncertainty is required to explain arc responses. Determining how probability information, or something akin to it, is represented in memory and used in decisions will require future work. Regardless of how the WM representation is structured, our findings highlight that current models of working memory must be consistent with the idea working memory representations contain not only an estimate of the stimulus, but a measure of memory uncertainty which can be used in subsequent decisions.

This work highlights how studying WM as a system integrated with decision-making can yield new insights on the capacity and representational nature of WM. By requiring participants to reason about memory contents, and manipulating the conditions under which these decisions were made, we show that WM representations contain a trial-level representation of probabilistic information, which is incorporated into subsequent decisions. This approach contrasts with that of many WM paradigms, which aim to minimize decision elements in order to examine WM in isolation (i.e. delayed estimation). However, there is no such thing as a “pure” working memory task – even simple paradigms involve reasoning about stored information. Furthermore, not only does WM involve decision-making, decisions in the real world often involve WM information (i.e. crossing the street, picking up objects). This has been recognized in reinforcement learning and behavioral economics, where task-relevant WM processes contribute to sequential value-based decisions^57–61^, risky decisions^62;63^, and delay discounting^64;65^. Thus, we suggest that models and theory should focus on understanding memory decision stages instead of minimizing their contribution. Studying WM in tasks with more decision elements could help us understand how the WM system functions realistically in parallel with other systems. Rather than treating WM storage capacity and decision-making as separate fields of inquiry, we suggest that an attempt to bridge these fields together will be necessary to understand working memory as a full and integrated system.

## Methods

### Experiment

#### Participants

12 people (7 female) participated in four 40-60 minute sessions of this experiment. Participants were aged 19-28 (mean 21.75) and recruited using flyers posted around New York University. All participants provided informed consent. This study conformed to the Declaration of Helsinki and was approved by the New York University Committee on Activities Involving Human Subjects.

#### Stimuli

The experiment was performed on a 40.9×32.5 cm Dell 1907FPC LCD monitor (1290×1025 resolution, refresh rate 75Hz) with a viewing distance of 40 cm. Participants were shown colored dots (1.26 degree radius) of eccentricity 6.3 degrees. Dot colors were sampled, with replacement, from a color space consisting of 360 equidistant colors using the cie L*a*b (centered at *L* = 54, *a* = 18, *b* = 8, with a radius of 59, available in MemToolbox^66^. Stimulus luminance was roughly 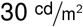 and background luminance was roughly 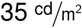. Colors were sampled with equal probability (uniform distribution) in sessions one and four and were drawn from a von Mises distribution with a fixed concentration parameter of *κ_w_* = 1.422 (60 degrees of circular standard deviation) and a randomly chosen mean for each participant in sessions two and three.

#### Trial procedure

We used a variant of delayed color estimation^6–8^. The participant was shown four colored dots for 100ms and, after a delay of 1000 ms, was asked to use the mouse to report on a color wheel the color of a randomly probed dot. After reporting an estimate, participants also made a judgment about the uncertainty in this response (Figure 1a), with a 1000 ms interval between trials.

#### Memory uncertainty report

We obtained a continuous report of memory uncertainty by asking participants to adjust the size of a symmetric “arc”, similar to a confidence interval, around their estimate to obtain reward (similar to Graf et al.^20^). If the true color was within the bounds of their arc, the participant received points inversely and linearly related to the size of the arc with the function 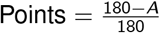, where *A* represents the size in degrees of half of the symmetric arc. If the true color was outside the bounds of the arc, they received 0 points. With this scoring rule, to get more points, participants had to use information about their memory quality to set an appropriate arc size, allowing us to use the arc size as a continuous metric of trial-by-trial memory uncertainty.

#### Performance incentivization

To incentivize participants to report memory uncertainty accurately, we attached time and monetary reward to the points obtained. Participants were awarded points on a 0-100 scale and permitted to leave when they achieved 18000 points total (or after an hour had elapsed). Participants completed an average of 1206 (904-1461) trials total with a mean of 645 (556-795) uniform stimulus distribution trials and 561 (348-645) von Mises stimulus distribution trials. Additionally participants received a performance bonus based on the summed points from three random trials (0-3 dollars). This incentive scheme is common in behavioral economics^67–69^ and encourages participants to evaluate each trial independently.

#### Stimulus distribution training, reinforcement, and testing

Participants were given extensive training to learn the stimulus distribution of each session. Participants were shown a movie consisting of 1000 samples of stimuli drawn from the session’s distribution. To provide a further visualization, samples were used to build a histogram of the session’s color distribution (Figure 1b). In von Mises sessions, during training participants further performed a “match assessment” (Figure 1b), where they used the mouse to simultaneously adjust the mean (left-right mouse position) and spread (up-down mouse position) of a sample of colored dots shown on screen to be representative of the learned distribution. Participants had to repeat this “match” assessment until they created a dot distribution close to the true distribution 3 consecutive times (mean error < 20 degrees, circular standard deviation error < 20 degrees).

Throughout the study, every 75 trials, participants were given a set of six binary choice questions about the session’s color frequencies to reinforce knowledge of the stimulus distribution (Figure 1b). Participants saw a bisected screen of two sets of dots and had to click on the set which best reflected the session’s color distribution. In order to continue the experiment, participants had to answer all six of these questions correctly. Participants received feedback and were given unlimited attempts to complete the questions. The foil distributions were slightly altered by shifting the mean color 60 degrees clockwise or counterclockwise (randomly) and the standard deviation by 30 degrees up or down in six combinations (narrow, wide, uniform, shifted, narrow-shifted, wide-shifted). The order of these foils was randomized in the questions.

To check whether participants learned the color frequencies, at the end of von Mises sessions, we had them report about their prior beliefs by generating a set of dots that matched the color distribution (similar to the earlier “match” assessment). They did this by using the mouse (left-right) to first select the most frequent color (mean of the distribution), and then moving the mouse (up-down) to set the spread of colors (circular standard deviation) (Figure 1b). Participants made three consecutive judgments and were awarded a performance bonus of 1 dollar if they reported the color frequencies accurately at least once (mean error < 20 degrees, circular standard deviation error < 20 degrees).

### Statistics

Unless otherwise specified, degrees of freedom is 11. Wilcoxon signed-rank tests are two-tailed. Correlations are Spearman correlations reported as the mean and s.e.m. across participants. For hypothesis testing on correlation coefficients, we use two-tailed t-tests on the Fisher-transformed coefficients. Bootstrapped confidence intervals are performed using 10000 samples with the bias corrected and accelerated percentile method^70^.

#### Confounding variable analysis

We tested whether the observed correlation between absolute error and arc size (*r*_obs_) was caused by both variables being correlated with a confounding variable such as stimulus color. To do this, we simulated data under the null hypothesis that the correlation is entirely caused by the stimulus color through shuffling our data. We sorted trials by stimulus color into groups of 5 trials with equal or similar stimulus color. Within each group, we then shuffled the arc size values uniquely such that each arc size was now associated with a different absolute error than on the trial it came from, but a similar stimulus color. This created a dataset where the relationship between arc size and error was solely dependent on the value of the stimulus color. We then correlated the shuffled arc sizes with the absolute error and obtained a value of *r*_null_ (similar to a semi-partial correlation). We repeated this 10,000 times for each participant to obtain the distribution of *r*_null_ and used this distribution to compute the probability of the observed *r*_obs_ per participant as well as the mean 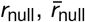. In addition to stimulus color, we apply the exact same analysis to the confounding variables of the circular variance of the four displayed items on each trial, and absolute color distance from the stimulus to the most confusable (closest in color space) item.

### Models

#### Base model

We model this task using a Bayesian decision model built on top of a variable-precision encoding model. We assume that on each trial, the observer encodes the stimulus *s* as a noisy memory *x* (Figure 3). We assume that *x* follows a von Mises distribution with mean *s* and precision (concentration) parameter *κ*,

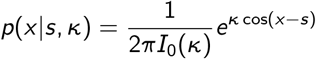

Variable precision means that *κ* itself varies from trial to trial; following previous work^9;39;40^, we assume it obeys a gamma distribution parameterized by its mean 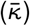 and shape parameter (*φ*),

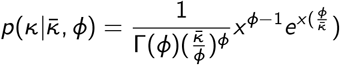

The model observer stores on each trial a representation of this precision *κ* as certainty *κ*\*, where we assume for the moment that *κ*\* = *κ*. The observer combines the memory *x* and associated memory uncertainty *κ*\* (von Mises) with their prior beliefs (von Mises) using Bayes’ rule to compute a posterior *p*(*s*|*x*). Since the product of two von Mises distributions is again a von Mises distribution, the posterior is a von Mises distribution with an analytically defined mean *μ*_pos_ and concentration *κ*_pos_^71^:

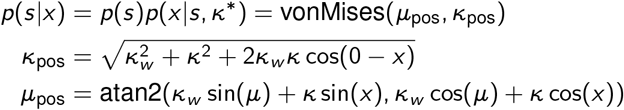

Where *μ* and *κ_w_* represent the mean and concentration parameter of the prior and *x* and *κ* represent the mean and concentration parameter of the likelihood function. We assume that the observer bases their prior beliefs on the learned stimulus distribution in the current session. Thus, we assume that in sessions 1 and 4, the prior was a uniform distribution, while in sessions 2 and 3, it was a von Mises distribution centered at the stimulus distribution mean; we allow its concentration parameter to be a free parameter, *κ_w_*.

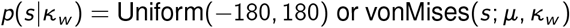

To obtain an estimate of the remembered color, 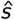, the observer samples from the posterior raised to a power *ν*, which represents decision noise^41^.

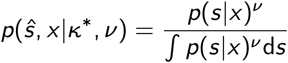

We assume that the probability that the the observer sets arc size *A* is a softmax function of the expected utility, with inverse temperature *β*,

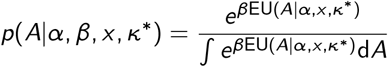

The expected utility is computed by multiplying the reward utility of an arc, *U*(hit), by the probability of getting that reward, *p*(hit).

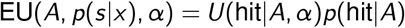

We assume that the utility of a correct response is not equal to the amount of points obtained (*R*(*A*), a linear function), but the points transformed by raised to a power *α*, representing risk attitude^42^.

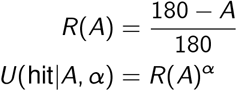

The probability of a hit is the integral of the posterior under the area covered by the arc. This is a von Mises distribution with parameters *μ*_pos_ and *κ*_pos_

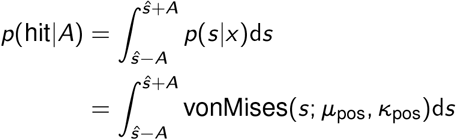

This is computationally demanding, since *p*(hit) depends on the stimulus estimate 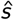, arc size *A*, mean of the posterior *μ*_pos_, and width of the posterior *κ*_pos_. To simplify, we can rewrite this expression, from which it is apparent that *p*(hit) is not a function of 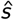 and *μ*_pos_, but the difference between them 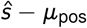

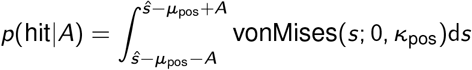

If we make the approximation that 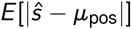 is small 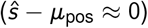:

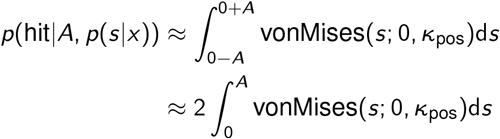

This removes the estimate from the integral and causing *p*(hit) to be only dependent on *A* and *κ*_pos_. This approximation has the effect of mildly underestimating arc sizes (see Supplementary material). However, since even after approximation, computing the von Mises integrals is computationally heavy we speed this up by precomputing a table of *p*(hit) for 180 *A* values and 1000 *κ*_pos_ values and linearly interpolating.

Furthermore, a lapse process is implemented such that on every trial there is a probability *λ* that the participant will have no information about the stimulus (for example due to a failure to encode the stimulus^72^). In this situation the observer will report a random estimate and an arc size according to the expected utility function associated with a flat posterior.

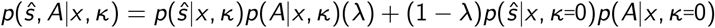

For the joint probability of 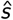 and *A* given the stimulus *s* and parameters 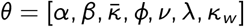, we obtain

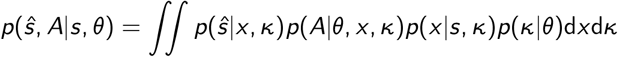

#### Modifications of the model

To test the assumptions of this model, we modify the base model (**vp-known** / **vp-k**, 6 parameters) in several ways:

- **VP-known (6p)**: No modifications Base model, a model in which the observer encodes stimuli with variable encoding precision (parameter *κ*) and represents this encoding precision as memory uncertainty (*κ*\*).
- **FP-known (5p)**: No variable precision A model in which people do not have internal fluctuations encoding precision, but always encode stimuli with a single fixed precision (*κ* = *κ*_fixed_). In this model, knowledge of memory quality is limited to knowledge of zero-precision trials (“lapses”).
- **VP-inaccessible (7p)**: No knowledge of memory uncertainty A model in which people have internal fluctuations in encoding precision, but have no knowledge of these fluctuations and assume their memory uncertainty is a fixed value (*κ*\* = *κ*_assumed_). In this model knowledge of memory quality is limited to knowledge of zero-precision trials (“lapses”). *κ*_assumed_ is a free parameter.
- **VP-discrete access (8p)**: Limited knowledge of memory uncertainty A model in which people have limited knowledge of their memory uncertainty: when it is above a threshold *κ*_threshold_ they assume it is a fixed value *κ*_assumed_, while if it is below the threshold they assume it is zero. *κ*_assumed_ and *κ*_threshold_ are free parameters.

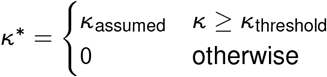

When the stimulus distribution is von Mises, to test how people incorporate prior and memory uncertainty information we test several different mechanisms of prior combination. Since an assessment of people’s prior knowledge revealed misconceptions about its width, the prior width in these models is a free parameter *κ_w_* unless otherwise stated. Models are named by their use of the prior during the task (first letter) or when lapsing (second letter) represented by the letters “Y” (*yes* prior use, prior width is a free parameter *κ_w_*), “N” (*no* prior use), or “T” (using the *true* stimulus distribution).

- **TT (6p)**: Always use the prior, perfect knowledge of prior Memory and prior are combined with Bayes’ rule. Prior width is not a free parameter but is fixed at the actual value (*κ_w_* = 1.422).
- **YY (7p)**: Always use the prior Memory and prior are combined with Bayes’ rule.
- **YN (7p)**: Use the prior, except when precision is zero Memory and prior are combined with Bayes’ rule, except when precision = 0. In that case, the observer does not use the prior and responds with a random estimate and an arc size based on a uniform posterior (random guess).
- **NY (7p)**: Only use the prior when precision is zero When responding from memory, participants ignore the prior. However, when precision = 0, the observer uses the prior to make an informed guess of the estimate, and sets an arc size based on the width of the prior.
- **NN (6p)**: Never use the prior or has a flat prior The prior is never used (*κ_w_* = 0).

#### Implementation

To evaluate the above integrals, we define a grid of 180 by 200 *x* and *κ* values in the respective ranges of [‒180,180] and [0,200] in degrees (in the *κ* dimension taking the integral in spacing with degrees of circular standard deviation instead of concentration units) and take a trapezoidal integral over both dimensions. Scaling up or down the resolution of these integrals does not change the results (see Supplementary material). Models are coded in MATLAB with tools from `circ_toolbox`^73^ and `mem_toolbox`^66^.

### Fitting

Each model has 5-9 free parameters which are fit for each participant using maximum-likelihood estimation, optimized through 50 randomly started runs of the Bayesian Adaptive Direct Search (`BADS`) algorithm^74^.

#### Model comparison

We compared models using k-fold (k=10) cross-validated log-likelihoods (LLcv), summed across folds. K-fold cross validation estimates the predictive power of a model while avoiding overfitting^75^. This is done by splitting the data into K folds, one of which is reserved for testing while the model is fit on the rest, this is then repeated for each fold. We take the summed log-likelihood of all of the testing folds as a measure of the model evidence. To combine model performance across participants, we average the per-trial LLcv and then multiply by the mean number of trials (across participants). We do so because the amount of trials varies across participants and is inversely correlated with their performance. Therefore, averaging individual-participant log-likelihoods would weigh poorly performing participants more (since those participants completed more trials). We also confirm our results with Akaike information criterion (AIC)^76^ and Bayesian information criterion (BIC)^77^ (see Supplementary material).

#### Fixed/Random odds ratio

To check whether there is evidence for participants using different models, we calculate the ratio of the probability of all participants following the same model (H0, fixed effects) against the probability of participants following different models (H1, random effects) given the data *y* and models *m*_1_…*m_K_* where *K* is the number of models. *p*(*y_n_*|*m_k_*) is the likelihood of the data for a participant *n* given a model *m_k_*.

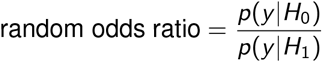

*p*(*y*|*H*_0_) is the probability of data being from a single model *m_k_*, multiplied by the prior probability of that model and summed for all possible models

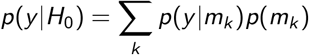

The probability of data *y* being from a model *m_k_* is the product of the model likelihood *p*(*y_n_*|*m_k_*) for each participant *n*, which is the sum of the log-likelihoods.

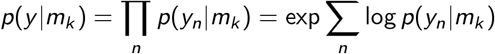

Assuming a uniform prior over models:

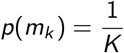

This gives us the full equation for *p*(*y*|*H*_0_):

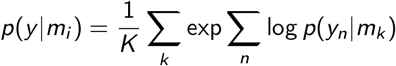

For *p*(*m*|*H*_1_), we use the probability of the random-effects group BMS model (see Rigoux et al., 2014^44^):

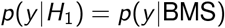

The fixed/random odds ratio reflects the relative probabilities of *H*_0_ and *H*_1_, large values reflect model evidence consistent with participants all being from the same model while small values are evidence that participants use different models.

## Acknowledgements

Thanks to Luigi Acerbi and Bas van Opheusden respectively for help with the random effects and shuffling analyses. Thanks to Aspen Yoo, Andra Mihali and Yanli Zhou for the useful conversations. This work was funded by grant number R01EY020958 to W.J.M.

## Author contributions

M.H., W.J.M., and D.F. designed the experiments. M.H. conducted the experiments. M.H. and W.J.M. developed the models. M.H. performed data analysis, visualization, and model fitting. M.H., W.J.M., and D.F. wrote the paper.

